# MULTIMAP: Multilingual picture naming test for mapping eloquent areas during awake surgeries

**DOI:** 10.1101/2020.02.20.957282

**Authors:** Sandra Gisbert-Muñoz, Ileana Quiñones, Lucia Amoruso, Polina Timofeeva, Shuang Geng, Sami Boudelaa, Iñigo Pomposo, Santiago Gil-Robles, Manuel Carreiras

## Abstract

Picture naming tasks are currently the gold standard for identifying and preserving language-related areas during awake brain surgery. With multilingual populations increasing worldwide, patients frequently need to be tested in more than one language. There is still no reliable testing instrument, as the available batteries have been developed for specific languages. Heterogeneity in the selection criteria for stimuli leads to differences, for example, in the size, color, image quality, and even names associated with pictures, making direct cross-linguistic comparisons difficult. Here we present MULTIMAP, a new multilingual picture naming test for mapping eloquent areas during awake brain surgery. Recognizing that the distinction between nouns and verbs is necessary for detailed and precise language mapping, MULTIMAP consists of a database of 218 standardized color pictures representing both objects and actions. These images have been tested for name agreement with speakers of Spanish, Basque, Catalan, Italian, French, English, Mandarin Chinese, and Arabic, and have been controlled for relevant linguistic features in cross-language combinations. The MULTIMAP test for objects and verbs represents an alternative to the DO 80 monolingual pictorial set currently used in language mapping, providing an open-source, standardized set of up-to-date pictures, where relevant linguistic variables across several languages have been taken into account in picture creation and selection.

## Introduction

Human language is a complex system of communication that supports the decoding, encoding, and transfer of information between individuals. It is a system that allows for communication not only about the here and now, but also about the past, the future, truths, lies, hopes, and desires. It is important for personal growth and socialization, but also for human development as the vehicle for cultural transmission. Losing or having impaired language ability can be a traumatic event that incurs hardship for affected individuals and those around them. Brain surgery procedures can unintentionally damage the language substrate, inducing impairments that can be irreversible (Duffau et al., 2005). For this reason, a patient is tested to identify eloquent areas that should not be removed in order to preserve or, in some cases, even improve, their quality of life (Ilmberger et al., 2008).

Since the late 1970s, when Ojemann and Mateer first reported using a visual object naming test during cortical stimulation in awake surgery (Ojemann & Mateer, 1979), the technique has become the gold standard in testing brain lesions involving language-related areas (De Witte & Mariën, 2013; Miceli, Capasso, Monti, Santini, & Talacchi, 2012). This language mapping procedure allows for the assessment of different language-related operations (i.e., access, retrieval and production of lexical-semantic information); the patient, presented with a series of drawings or pictures, is asked to name the depicted objects using a noun.

Although the use of object naming tasks is widespread across surgical teams in many different geographical locations, heterogeneity in the stimuli selection criteria of previous batteries (i.e., differences in picture size, color, image quality, name agreement), in addition to the use of morphologically and typologically different languages across different studies (see supplementary material for a table review), has greatly hindered the comparison and generalization of results. To our knowledge, a battery allowing for direct comparison between two languages in awake surgery has never been designed, even if some teams have tested multilingual patients in more than one language (Giussani, Roux, Lubrano, Gaini, & Bello, 2007). The critical need for a multilingual approach is evident given reports on bilingual aphasia where, in some cases, patients proficient in two languages prior to the lesion are selectively impaired in one after surgery (Fabbro, 2001). It is also known from the aphasic literature that linguistic variables, such as word frequency and imageability, among others, influence lexical performance (Luzzatti et al., 2002). However, multilingual brain stimulation studies have not reported how they controlled stimuli across languages, despite the fact that this choice of stimuli could influence findings concerning the brain areas that are common or specific to these languages (Bello & Acerbi, 2006; Cervenka, Boatman-Reich, Ward, Franaszczuk, & Crone, 2011; Roux, Lauwers-Cances, Trémoulet, Mascott, & Démonet, 2004). For these reasons, and given that the multilingual population in our society continues to increase as more and more people know and employ a second language in their daily lives, a multilingual evaluation tool has become necessary (EuroStat, 2015). In this paper, we present MULTIMAP, a new multilingual battery of standardized pictures in Spanish, with norms for Basque, Catalan, Italian, French, English, Mandarin Chinese, and Arabic. This is a tool that will allow surgical teams to test patients in bilingual contexts in a controlled and comparable manner.

We first conducted a systematic revision of the literature on naming tasks for awake surgery (see supplementary material for a table review). Out of the 52 articles that reported using an object naming task, 10 included an introductory sentence (e.g. “This is…”) printed above the picture to elicit the production of a determiner-noun pair (e.g. “This is… an apple” instead of “apple”) (Hamberger et al., 2016; Hamberger, Seidel, Goodman, & McKhann, 2010; Ille et al., 2015; Khan, Herbet, Moritz-Gasser, & Duffau, 2014; Lubrano, Roux, & Démonet, 2004; Moritz-Gasser & Duffau, 2009; G. Ojemann & Mateer, 1979; Roux, Borsa, & Démonet, 2009; Roux, Boukhatem, Draper, Sacko, & Démonet, 2009; Rutten, 2015). Patients are required to overtly produce grammatical information related to the selection of the appropriate determiner, as it encodes number and, in some languages, also gender information (i.e., in Spanish “una^f.sg.^ manzan<u>a</u> ^f.sg.^” [“an apple”]). This kind of task allows for the identification of different types of errors, most frequently: speech arrest, in which the patient is unable to speak; and anomia, where the patient can read the introductory phrase, but cannot retrieve and produce the noun^1^.

In addition to the object naming task, some neurosurgery teams have introduced verb tests in their practices. We retrieved 13 studies that reported the use of images to elicit verb production (Chen, Tan, Deng, & Xu, 2010; Conner, Chen, Pieters, & Tandon, 2014; Corina et al., 2005; Havas et al., 2015; Herholz et al., 1997; Lubrano, Filleron, Démonet, & Roux, 2014; J. G. Ojemann, Ojemann, & Lettich, 2002b; Papagno et al., 2011; Rofes et al., 2017; Roux et al., 2003; Sierpowska et al., 2015; Skrap, Marin, Ius, Fabbro, & Tomasino, 2016; Tomasino et al., 2014). In these tasks, patients were presented with a drawing or a picture of an action and they were asked to produce the appropriate verb: either a finite or an infinitive form, depending on the specific requirements of the task. The generation of finite verbs, unlike infinitives, requires the production of inflectional features that, depending on the language, may encode for number, gender, person, and/or time. Verbs refer to events and imply the projection of a complex representation in which the agent is associated with a specific thematic role and they differ from nouns at the lexical, semantic, morphological, and syntactic levels. In addition, the number of morphologically inflected forms is higher for verbs than for nouns.

This distinction between nouns and verbs has been demonstrated at the behavioral, electrophysiological, and neuroanatomical levels (Vigliocco, Vinson, Druks, Barber, & Cappa, 2011), demonstrating a double dissociation that should be taken into account when planning an awake surgery. Notably, direct cortical stimulation studies also show this double dissociation when object and action naming tasks are used, and allow for the identification of distinct territories in frontal and temporal brain areas in which stimulation selectively impairs verb or noun production (Corina et al., 2005; Crepaldi, Berlingeri, Paulesu, & Luzzatti, 2011; Lubrano et al., 2014; J. G. Ojemann, Ojemann, & Lettich, 2002a; Rofes et al., 2017). From these studies, it seems clear that comparing nouns and verbs is necessary for a detailed and precise language mapping procedure.

Several instruments have been previously employed in this endeavor. The most common batteries reported for mapping nouns are the Oral Denomination 80 (Metz-Lutz, Kremin, & Deloche, 1991) (DO 80); the images included in the Boston Diagnostic Aphasia Examination (Goodglass & Kaplan, 1972) (BDAE); and the pictures from the Snodgrass and Vanderwaart battery (Snodgrass & Vanderwart, 1980). These sets of images were not designed for intraoperative language mapping, and name agreement norms and values for relevant linguistic variables, such as word frequency, length, or familiarity, are not provided or are only available for some of the languages in which they have been employed. In addition, payment is required to access both the DO 80 and the BDAE, restricting their use. In some other studies using intraoperative object naming, stimuli are only described as “black-and-white drawings”, “simple drawings” or just “pictures representing common objects”, and no further information is given about their construction or the linguistic variables of the target words. In studies that have carried out language mapping using verbs, some teams have reported using the same pictures for verb action naming that they use in their object naming tasks (Papagno et al., 2011), while others have presented pictures depicting actions (Havas et al., 2015). For verbs, as for nouns, only some studies have reported controlling their stimuli for the relevant linguistic variables or have described how these stimuli were constructed and selected.

To address these shortcomings and the critical demand for multilingual material, we developed a multilingual picture naming test (MULTIMAP) for the mapping of eloquent areas during awake brain surgery. MULTIMAP consists of a database of standardized color pictures of common objects and actions, tested for name agreement measures in speakers of Spanish, Basque, Catalan, Italian, French, English, Mandarin Chinese, and Arabic. In the MULTIMAP object and action naming subtests, items were independently selected for each language and controlled for a number of linguistic variables (i.e. frequency, length, and familiarity). We also equated variables for Spanish and each of the other languages separately to facilitate testing in multilingual patients. This new battery will help teams plan surgical interventions, providing them with a sensitive, validated instrument for intraoperative language mapping offering better results in terms of the patient’s health and overall quality of life. In the following sections, we include a detailed description of the material and its validation in order to facilitate replication and extension to other languages in future research.

## Material and Procedure

### Participants

One hundred and twenty-three healthy adult volunteers aged 18 to 60 years were recruited and paid for the object naming task, and 124, within the same age range, for the verb naming task. A detailed description of the sample used for each language is included in Table 1 (i.e., sample size, ages, gender, educational level, and linguistic profile). They all had normal or corrected to normal vision and no history of psychiatric, neurological diseases, or learning disabilities. A signed informed consent was collected from each participant as stipulated in the ethics approval procedure of the BCBL^2^ Research Ethics Committee. All of the participants were from the Basque Country in Spain and were highly proficient Spanish-Basque bilinguals.

**Table 1:** Description of the participants for the three phases of the study. N=number of participants for each of the tasks in each of the phases.

|  | Task language | Task | N | Gender | Mean Age (SD) |
| --- | --- | --- | --- | --- | --- |
| **Phase 1:**<br>**Material Selection** | Spanish | Objects | 123 | Female = 98 | 39.98 (9.30) |
|  |  | Verbs | 124 | Female = 101 | 39.96 (9.49) |
| **Phase2:**<br>**Validation** | Spanish | Objects | 20 | Female = 13 | 35.35 (13.64) |
|  |  | Verbs | 20 | Female = 13 | 35.35 (13.64) |
| **Phase 3:**<br>**Cross-language combinations** | Basque | Objects | 100 | Female = 82 | 39.66 (9.98) |
|  |  | Verbs | 95 | Female = 76 | 39.24 (10.34) |
|  | Catalan | Objects | 100 | Female = 70 | 34.14 (11.13) |
|  |  | Verbs | 100 | Female = 70 | 34.14 (11.13) |
|  | Italian | Objects | 101 | Female = 71 | 32.09 (10.07) |
|  |  | Verbs | 101 | Female = 71 | 32.09 (10.07) |
|  | French | Objects | 104 | Female = 89 | 24.29 (9.85) |
|  |  | Verbs | 104 | Female = 89 | 24.29 (9.85) |
|  | English | Objects | 99 | Female = 75 | 35.42 (11.56) |
|  |  | Verbs | 99 | Female = 75 | 35.42 (11.56) |
|  | Chinese | Objects | 128 | Female = 87 | 23 (3.68) |
|  |  | Verbs | 128 | Female = 87 | 23 (3.68) |
|  | Arabic | Objects | 101 | Female = 73 | 25.05 (7.15) |
|  |  | Verbs | 101 | Female = 73 | 25.05 (7.15) |

### Material Selection and Norming

The development of the MULTIMAP test comprised three phases. **Phase 1** included the selection of the linguistic material and its pictorial representations, taking into account the following psycholinguistic variables for each target word: word frequency, number of letters, number of phonemes, number of syllables, number of substitution neighbors, and name agreement. In order to determine whether the final set of stimuli were sufficiently good to test patients during surgery, we included **Phase 2**. This stage involved the final validation of the selected material in a different group of typical participants and the standardization of the protocol as indicated in awake surgery guidelines (i.e., name agreement, presentation times, and accuracy per item). During **Phase 3**, we carried out a cross-language validation of the stimuli, controlling for word frequency, number of letters, number of phonemes, number of syllables, number of substitution neighbors, and name agreement for seven pairs of languages: Spanish-Basque, Spanish-Catalan, Spanish-Italian, Spanish-French, Spanish-English, Spanish-Mandarin Chinese, and Spanish-Arabic.

### Phase 1 (Material Selection)

We selected an initial list of 109 nouns and 109 action verbs from the EsPal database (http://www.bcbl.eu/databases/espal/)(Duchon, Perea, Sebastián-Gallés, Martí, & Carreiras, 2013). These words were selected to cover a wide range of frequencies and semantic fields. As the norming data was to be collected within the Basque Country, where most people are Spanish-Basque bilinguals, we avoided all Spanish-Basque cognates. Additionally, we excluded plural invariant forms (i.e. *tijeras*, scissors) and nouns and verbs that could have negative valence for the target audience of the battery (i.e. *cerebro*, brain; *morir*, to die). All the selected verbs were transitive and non-reflexive. A graphic designer was commissioned to create the initial set of color drawings using the same dimensions (988 x 719 pixels; 8.37 x 6.09 centimeters; DPI = 300 pixels/centimeter) and similar styles for the whole set. Verbs were depicted with a human agent (see Figure 1 for an example).

**Figure 1:**
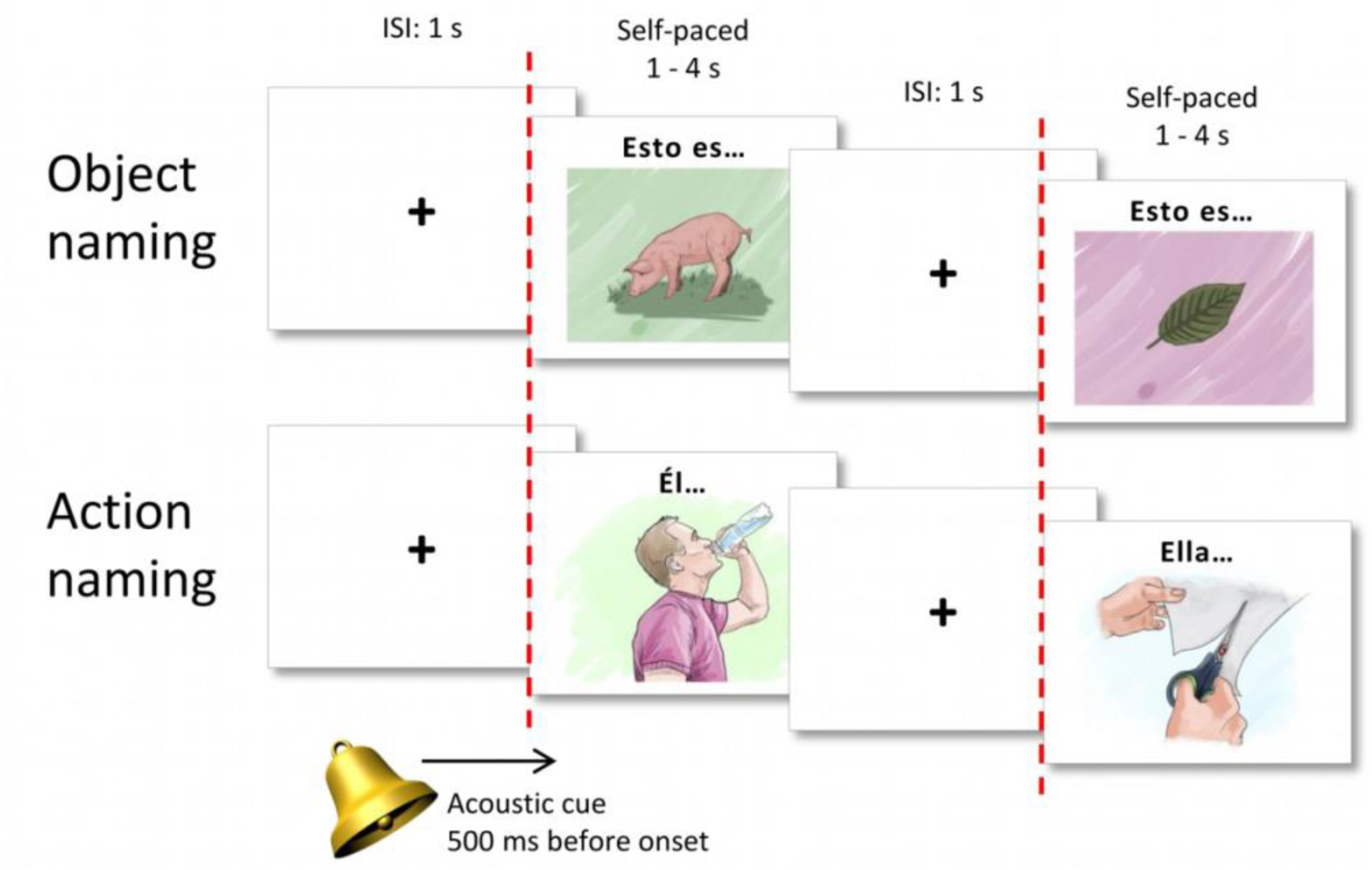
Example of stimulus presentation for both object and action naming. An acoustic cue is presented during the fixation cross, 500ms before the stimulus onset. The fixation cross appears on the screen for 1s, followed by the target picture which appears for 1 to 4s, self-paced as determined by the speed of the patient’s response.

The name agreement data was collected using the Lime Survey platform (https://www.limesurvey.org/es/) in two separate survey sections: nouns and verbs. Participants were contacted by email with a detailed explanation of the study, and instructions were provided before the start of each part. For both nouns and verbs, the instructions indicated that participants would see a set of images presented one at a time and should write just one word to name each picture: a noun for the objects and an infinitive verb for the actions. If participants were not sure what the drawing represented or did not know the label for it, they were instructed to write “NS” for “no sé” [“I don’t know”, in Spanish]. The drawings were randomly presented one by one in the center of a computer screen. The experimental session was self-paced and lasted about 30 minutes.

A final subset of 88 drawings including 44 objects and 44 actions, with a name agreement of at least 80%, was selected. There were no significant differences in frequency, number of letters, number of phonemes, number of syllables, number of substitution neighbors, or familiarity in picture names. Values for imageability and concreteness were high for both nouns (mean imageability = 6.20, SD = 0.37; mean concreteness = 5.88, SD = 0.47) and verbs (mean imageability = 5.25, SD = 0.57; mean concreteness = 4.73, SD = 0.65), but could not be equated given the semantic characteristics of verbs.

### Phase 2 (Validation)

Once we had the final subset of 88 images, we prepared the images for use in the awake surgery setting. Above each object, we added the text “Esto es…” [“This is…” in Spanish] to force participants to produce a short sentence that would have to agree in number and gender with the target noun. Above the action pictures, we included a noun phrase to be used as the subject of the sentence, that is either “Él…” or “Ella……” [“He…” or “She…” in Spanish] depending on the gender of the agent. This introductory text was used as a cue for the production of a sentence that started with the given subject and had a finite verb form in 3^rd^ person singular. We used MATLAB version 2012b and Cogent Toolbox (http://www.vislab.ucl.ac.uk/cogent.php) to present the images, as they would be used in a surgery setting. First, a white screen with a black fixation cross appears for 1s, followed by a picture presented for 4s. An acoustic cue is given 500ms before the onset of each stimulus during the fixation cross (the Matlab script and its compiled version are available for use at https://git.bcbl.eu/sgisbert/multimap2).

### Phase 3 (Cross-language combinations)

Seven norming studies were carried out. The first one included the Spanish set of stimuli, described in Phase 1, and the other five comprised the different cross-language combinations (i.e., Spanish-Basque, Spanish-Catalan, Spanish-Italian, Spanish-French, and Spanish-English). For each of these combinations, pictures were tested for name agreement, following the same procedure described in Phase 1, in samples of highly proficient users of the languages. Frequency, length, and information about orthographic neighbors were extracted for the words in each language from the following databases: for Basque, e-Hitz(Perea et al., 2006); for Catalan, the Corpus Textual Informatitzat de la Llengua Catalana (https://ctilc.iec.cat/), accessed through NIM(Guasch, Boada, Ferré, & Sánchez-Casas, 2013) (http://psico.fcep.urv.es/utilitats/nim/); for Italian, CoLFIS(Bambini & Marco, 2012) (http://linguistica.sns.it/esploracolfis/home.htm); for French, Lexique (http://www.lexique.org/); for English, the British National Corpus (http://www.natcorp.ox.ac.uk/), accessed through NIM(Guasch et al., 2013) (http://psico.fcep.urv.es/utilitats/nim/) and the Glasgow Norms(Scott, Keitel, Becirspahic, Yao, & Sereno, 2019); for Mandarin Chinese, SUBTLEX-CH (Cai & Brysbaert, 2010) (http://crr.ugent.be/programs-data/subtitle-frequencies/subtlex-ch); and for Arabic, Aralex (Boudelaa & Marslen-Wilson, 2010). Stimuli with at least 80% name agreement were selected so that nouns and verbs did not show significant differences in frequency, number of letters, picture-name agreement, and H-index, both within and between the two languages of each pair.

Depending on the language, sentence production requires the use of different grammatical devices (e.g., case markings, word order, inflectional morphology, etc.). Thus, in addition to orthographical and lexical factors, some syntactic constraints were applied. For all languages, we took out plural invariant nouns (i.e. “Tijeras”, scissors) and kept only transitive verbs. For each of the cross-language combinations, we removed cognates.

### Data processing

A native speaker of each language checked the answers for writing/spelling errors, standardized the writing using capital letters, and merged basic variants of the same target word (e.g. hyphenated, pluralized forms). After that, we excluded trials where participants did not know the name or did not recognize the concept from all analyses. Name agreement and H-index(Snodgrass & Vanderwart, 1980) were then computed for the remaining 109 nouns and 109 verbs. Both of these measures reflect the level of agreement across participants: while name agreement represents the percentage of participants who give a certain answer, the H-index reflects response variability in terms of the number of different answers given by participants.

Given the name agreement data for each language, we extracted items with a score of at least 80%. With these items, we first created a list for each language, selecting the same number of nouns and verbs, and ensuring that there were no significant differences across the linguistic values we controlled for (i.e. frequency, length, orthographic neighbors) using unpaired two-sample T tests. Next, we paired each of the languages with Spanish to create the bilingual tasks. Here, we selected new items from the 80% name agreement list to create lists of the same length between languages, again controlled for frequency, H-index, length, and orthographic neighbors (except for the combinations Spanish-Mandarin Chinese, were length and orthographic neighbors were not contemplated, and Spanish-Arabic, where orthographic neighbors were not contemplated). We calculated ANOVAs between the four lists (objects language 1 x objects language 2 x verbs language 1 x verbs language 2) to check that there were no significant differences in the relevant variables.

## Results

### Phase 1 (Material Selection)

We tested 109 pictures of objects and 109 pictures of actions in a Spanish speaking sample. From this picture pool with a constraint of at least 80% name agreement, we selected 44 objects and 44 verbs that showed no significant differences in name agreement, frequency per million, number of letters, number of phonemes, number of syllables, and familiarity (see Table 2 for an overview of the variables and t-test results comparing objects and verbs within languages). The values for imageability and concreteness showed statistically significant differences, with objects higher than verbs in both cases. Imageability can be defined as the ease with which a word brings to mind a sensory image, while concreteness is the property of being able to see, hear and touch something (Bird, Howard, & Franklin, 2003). Verbs as a class are by definition less imageable and concrete than objects, given that objects that can be drawn have stronger sensory features than action verbs, which are primarily functional and motoric(Bird, Howard, & Franklin, 2000).

**Table 2:**
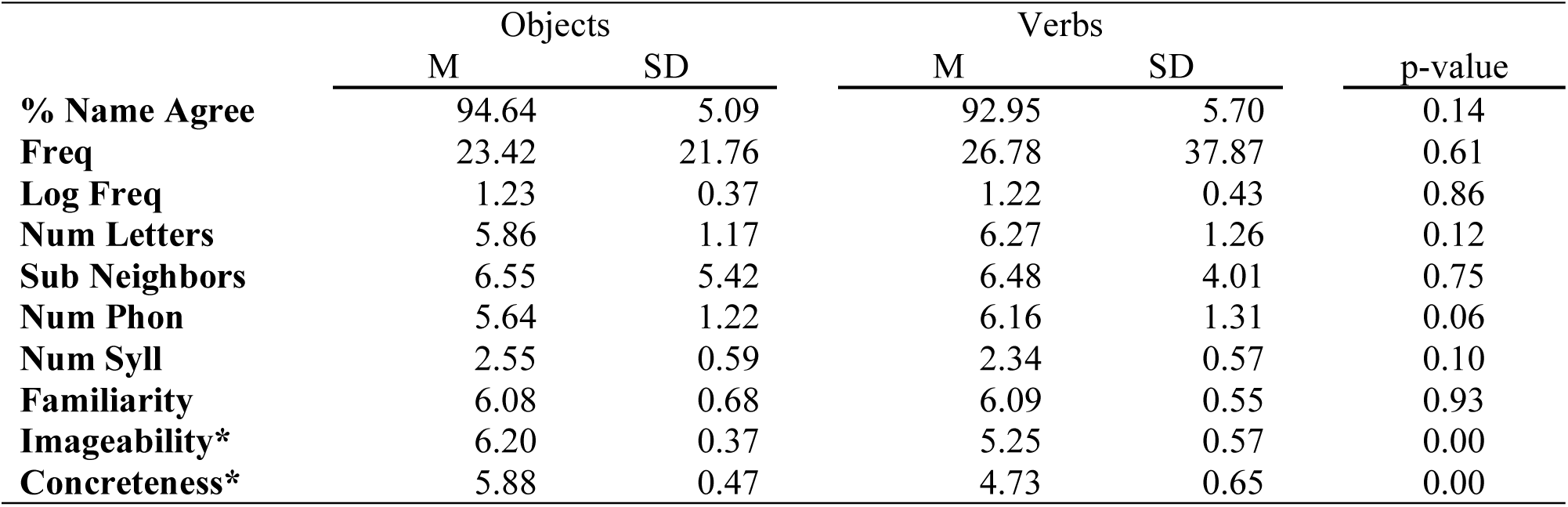
Mean values and standard deviations of the norming variables for nouns and verbs. Percentage of name agreement was calculated from the participants’ answers. Frequency per million, logarithmic frequency, number of letters, number of substitution neighbors, number of phonemes, number of syllables, familiarity, imageability and concreteness values were taken from the EsPal database.

|  | Objects |  | Verbs |  | p-value |
| --- | --- | --- | --- | --- | --- |
|  | M | SD | M | SD |  |
| **% Name Agree** | 94.64 | 5.09 | 92.95 | 5.70 | 0.14 |
| **Freq** | 23.42 | 21.76 | 26.78 | 37.87 | 0.61 |
| **Log Freq** | 1.23 | 0.37 | 1.22 | 0.43 | 0.86 |
| **Num Letters** | 5.86 | 1.17 | 6.27 | 1.26 | 0.12 |
| **Sub Neighbors** | 6.55 | 5.42 | 6.48 | 4.01 | 0.75 |
| **Num Phon** | 5.64 | 1.22 | 6.16 | 1.31 | 0.06 |
| **Num Syll** | 2.55 | 0.59 | 2.34 | 0.57 | 0.10 |
| **Familiarity** | 6.08 | 0.68 | 6.09 | 0.55 | 0.93 |
| **Imageability\*** | 6.20 | 0.37 | 5.25 | 0.57 | 0.00 |
| **Concreteness\*** | 5.88 | 0.47 | 4.73 | 0.65 | 0.00 |

### Phase 2 (Validation)

The final subset of 88 images was validated in a group of 20 Spanish native speakers that performed the task as would be done in the surgical setting. The responses were recorded to measure the time needed to produce the whole sentence “This is…” + determiner^[masc/fem]^ + target noun^[masc/fem]^ or pronoun^[masc/fem]^ + verb ^[3rd person singular, masc/fem]^. The response times did not, in any case, reach the maximum 4s allotted (mean response time for objects = 1.39 s, SD = 0.31; mean response time for actions = 1 s; SD = 0.35).

### Phase 3 (Cross-language combinations)

For bilingual lists, objects and verbs in both languages were always equated for name agreement (higher than 80% in all cases) and controlled so that linguistic variables did not show significant differences (see Table 3 for an overview of the variables and t-test results comparing objects and verbs within languages). The resulting monolingual lists, with objects and verbs controlled, are all 40 items long, except for the Mandarin Chinese version, where lists are 31 items long and the Arabic version, with 30 items per list. The bilingual lists, which combine four lists (objects language 1 x objects language 2 x verbs language 1 x verbs language 2), are shorter and they vary in length. The Spanish-Basque combination lists are 30 items long (word frequency F = 1.59, p = 0.22); the Spanish-French, 25 items long (word frequency F = 0.52, p = 0.67); the Spanish-English, 30 items long (word frequency F = 1.13, p = 0.34); the Spanish-Mandarin Chinese lists have 25 items (word frequency F = 0.58, p = 0.63); and the Spanish-Arabic lists are 30 items long (word frequency F = 0.21, p = 0.89).

**Table 3:** Mean values and standard deviations of the norming variables for nouns and verbs in each of the seven languages. Percentage of name agreement and H-index were calculated from the participants’ answers. Frequency per million, logarithmic frequency, and number of letters values were taken from the databases listed in the Material Selection and Norming (Phase 3) section.

|  | Objects |  | Verbs |  | p-value |
| --- | --- | --- | --- | --- | --- |
|  | M | SD | M | SD |  |
| Spanish |  |  |  |  |  |
| % Name Agree | 94.19 | 5.77 | 94.68 | 4.37 | 0.67 |
| Freq | 20.65 | 17.18 | 22.39 | 19.28 | 0.88 |
| Log Freq | 1.22 | 0.32 | 1.21 | 0.40 | 0.21 |
| Num Letters | 5.80 | 1.45 | 6.18 | 1.17 | 0.56 |
| H-Index | 0.33 | 0.30 | 0.35 | 0.26 | 0.67 |
| Basque |  |  |  |  |  |
| % Name Agree | 93.66 | 5.51 | 92.75 | 6.26 | 0.49 |
| Freq | 94.93 | 105.62 | 99.63 | 81.18 | 0.83 |
| Log Freq | 1.82 | 0.38 | 1.73 | 0.61 | 0.41 |
| Num Letters | 6.75 | 1.46 | 6.30 | 1.78 | 0.22 |
| H-Index | 0.40 | 0.29 | 0.49 | 0.45 | 0.30 |
| Catalan |  |  |  |  |  |
| % Name Agree | 95.31 | 5.84 | 94.18 | 94.18 | 0.40 |
| Freq | 43.35 | 34.54 | 43.83 | 43.83 | 0.95 |
| Log Freq | 1.50 | 0.38 | 1.53 | 1.53 | 0.75 |
| Num Letters | 6.00 | 1.62 | 6.50 | 6.50 | 0.13 |
| H-Index | 0.28 | 0.28 | 0.34 | 0.34 | 0.26 |
| Italian |  |  |  |  |  |
| % Name Agree | 94.51 | 6.67 | 94.84 | 5.85 | 0.81 |
| Freq | 79.13 | 134.77 | 78.42 | 70.82 | 0.98 |
| Log Freq | 2.23 | 0.54 | 2.33 | 0.46 | 0.34 |
| Num Letters | 7.28 | 1.78 | 7.68 | 1.38 | 0.27 |
| H-Index | 0.30 | 0.33 | 0.31 | 0.33 | 0.85 |
| French |  |  |  |  |  |
| % Name Agree | 95.04 | 4.27 | 93.66 | 5.86 | 0.23 |
| Freq | 60.37 | 82.05 | 46.87 | 45.32 | 0.36 |
| Num Letters | 6.00 | 1.66 | 6.65 | 1.35 | 0.06 |
| H-Index | 0.29 | 0.21 | 0.40 | 0.33 | 0.07 |
| English |  |  |  |  |  |
| % Name Agree | 94.10 | 6.33 | 93.96 | 6.24 | 0.92 |
| Freq | 88.60 | 108.53 | 90.15 | 88.22 | 0.94 |
| Log Freq | 1.72 | 0.44 | 1.74 | 0.48 | 0.83 |
| Num Letters | 4.88 | 1.45 | 4.50 | 0.99 | 0.18 |
| H-Index | 0.35 | 0.32 | 0.33 | 0.30 | 0.78 |
| Chinese |  |  |  |  |  |
| % Name Agree | 94.80 | 5.67 | 92.87 | 6.84 | 0.23 |
| Freq | 118.06 | 157.27 | 119.23 | 193.77 | 0.98 |
| Log Freq | 1.72 | 0.6 | 1.66 | 0.7 | 0.71 |
| H-Index | 0.77 | 0.51 | 0.83 | 0.46 | 0.63 |
| Arabic |  |  |  |  |  |
| % Name Agree | 90.99 | 6.60 | 90.2 | 6.78 | 0.65 |
| Freq | 12.59 | 16.63 | 13.59 | 15.71 | 0.81 |
| Num Letters | 3.93 | 0.23 | 4.13 | 0.43 | 0.23 |
| H-Index | 0.53 | 0.36 | 0.59 | 0.39 | 0.53 |

## Discussion

Language is the main vehicle humans use to communicate and transfer information. Groups of neurons arranged in networks support language functions. Any modification to this system (e.g., brain lesions, epilepsy seizure, etc.) may irreversibly impair language capacity, leading to unexpected and problematic consequences for the affected individual. For this reason, brain surgeries are increasingly planned in an awake setting, making it possible to monitor patients’ language function and spare brain tissue that is indispensable (De Witte & Mariën, 2013). There is a critical need for specific tools, based on linguistic, neuroscientific, and clinical knowledge about how the brain decodes and encodes linguistic information, that can facilitate precise mapping of these eloquent regions during neurosurgery.

To address this need, we developed the MULTIMAP test, a multilingual picture naming task including both objects and actions for mapping eloquent areas during awake brain surgery. Images included in the MULTIMAP test are colored drawings of objects and actions that have been standardized in seven different languages (Spanish, Basque, Catalan, Italian, French, English, Mandarin Chinese, and Arabic), controlling for name agreement, frequency, length, and substitution neighbors. This image database was designed to minimize linguistic distance between different groups of items, allowing direct comparisons between objects and actions within and across languages. This new set of standardized pictures will be an important and useful tool, enabling neurosurgeons to intraoperatively map language functions while taking into account the double dissociation between nouns and verbs reported in the literature and thus increasing presurgical and surgical mapping sensitivity for the detection of active eloquent brain areas. In addition, these materials will improve language mapping in multilingual patients, facilitating the identification and preservation of areas that show interference in only one of their languages that would not be detected by a monolingual test. MULTIMAP thus offers two important improvements over other picture naming tasks reported in the literature: the inclusion of objects and actions and multilingual norming data.

In spite of empirical evidence demonstrating neuroanatomical distinctions for nouns and verbs (Vigliocco et al., 2011) and native and second/third languages (Giussani et al., 2007), there is no available material that tackles both of these issues at the same time. In this regard, MULTIMAP constitutes the first tool designed to explore these factors in a structured way, in order to identify and preserve eloquent brain tissue, and to obtain better results in terms of patients’ health and overall quality of life. MULTIMAP includes two separate tests, one for objects and one for actions to address the noun/verb issue; both tests are controlled so that there are no significant differences for linguistic variables such as frequency, length, and orthographic neighbors. The need to map both objects and actions is motivated by evidence of a double dissociation, demonstrated at the behavioral, electrophysiological, and neuroanatomical levels (Vigliocco et al., 2011) and, as pointed out in the Introduction, also reported in direct cortical stimulation studies. From the 52 reviewed studies where a picture naming task had been used in the context of awake brain surgery, we found only 13 that included a verb task in addition to object naming. Although these tasks were varied in their requirements and the nature of the stimuli they employed, they all identified distinct territories, mainly in frontal and temporal brain areas where stimulation impaired verb and noun production separately (Corina et al., 2005; Crepaldi et al., 2011; Lubrano et al., 2014; Ojemann et al., 2002a; Rofes et al., 2017). Moreover, our tasks include an extra level of complexity beyond the extraction of morphosyntactic information, as the production of target objects and actions have been embedded in simple sentences, such as “This is a house” for objects, and “He/she sings” in the case of actions. This entails a higher level of complexity since the generation of such sentences requires the projection of representations in which thematic roles are assigned to different elements in the sentence (i.e., He^agent^ sings), in addition to matching the target word to its real-world referent. Therefore, combining object and verb processing tasks at the sentence level ensures a more accurate and thorough mapping of language functions, helping to more accurately identify and preserve the neural linguistic substrate essential for a patient’s quality of life.

The second improvement offered by MULTIMAP is its multilingual nature. Neuroimaging studies in bilinguals suggest that there is a common cerebral organization across languages, but also describe activations specific to each of the languages (Kim, Relkin, Lee, & Hirsch, 1997; Marian, Spivey, & Hirsch, 2003; Rueckl et al., 2015). Direct electrostimulation studies have also revealed language-specific areas (Giussani et al., 2007).This study concluded that multilingual patients should be tested in all languages in which they are fluent during brain mapping procedures so as to avoid selective or preferential impairments. With this objective in mind, the images included in MULTIMAP were tested in seven languages taking into account the lexical and morphological features of each language. This resulted in seven separate sets of object and action pictures with at least 80% name agreement in their target language, accounting for relevant linguistic variables like frequency and word length. These sets can be combined in controlled bilingual sets, enabling researchers from different countries to use the same materials, and to compare results not only from monolingual samples but also in cross-linguistic research on multilingual patients. It will also play a role in the postoperative quality of life for multilingual patients, as these materials will facilitate the identification and preservation of areas where interference impairs only one of their languages (Giussani et al., 2007), areas that might not be detected by monolingual tests.

MULTIMAP represents an alternative to the pictorial sets like the DO 80 or the BDAE currently used in language mapping, and overcomes their limitations by providing a free, standardized battery of up-to-date pictures where relevant linguistic variables have been taken into account in creating and selecting sets for each of the presented languages. The test includes between 25 and 30 items, depending on the language or language combinations selected, and can be performed in a maximum of 5 minutes per language. This number of trials should allow surgeons to identify eloquent areas that generate errors at least two out of three times when stimulated, without exceeding the safe awake time for the patient. In compliance with standards for direct cortical stimulation which aim to minimize intraoperative risk for patients, all of the items have been tested so that the required answer can be produced in less than 4s. The complete set of stimuli and the norming data are available at (https://git.bcbl.eu/sgisbert/multimap2), including a Microsoft Excel spreadsheet with information on the initial 218 images. This spreadsheet contains values for each of the seven languages, including name agreement and other relevant linguistic variables (i.e. frequency, word length, and number of orthographic neighbors). It contains the filter information to create the seven individual object-verbs lists per language, and bilingual lists for the combinations Spanish-Basque, Spanish-French, Spanish-English, Spanish-Mandarin Chinese, and Spanish-Arabic. Other language combinations can be created with the information provided in the tables. The materials are open-access and free from copyright restrictions for non-commercial purposes.

## Supporting information

Literature Revision Table

## Acknowledgements

We would like to thank the BCBL Lab Department for data recordings and Magda Altman for her useful comments on the manuscript.

## Study Funded

by Severo Ochoa (SEV-2015-049); the European Research Council (ERC-2011-ADG-295362), and MINECO (RTI2018-093547-B-I00).

## Footnotes

1 Other errors that can be identified using this kind of task are (1) semantic paraphasia if, instead of the target noun, a semantically related word is produced (e.g. “tiger” for “lion”); (2) phonological paraphasia, when the patient produces the target word with phonological deviations (e.g. “fable” for “table”); (3) neologism creation, inventing new word; or (4) perseveration, if the patient repeats an item, even after a new image is presented. Other significant limitations can also be detected using this task, such as delays in producing a response and hesitations.

2 Basque Center on Cognition, Brain and Language

