## Supplementary material for "MULTIMAP: Multilingual picture naming test for mapping eloquent areas during awake surgeries": Literature Revision Table

**Title:** MULTIMAP: Multilingual visual naming test for the mapping of eloquent areas during awake surgeries.

Table 1: Summary of the literature review. We used the keywords *object, verb, naming, awake surgery, neurosurgery,* and *cortical stimulation* on PubMed to select studies where naming tasks using objects and/or verbs had been used during a brain awake surgery procedure.

| Study | Language | Tasks | Stimuli |
|---|---|---|---|
| (Ojemann & Mateer, 1979) | English[1] | Object picture naming → "This is…" | |
| (Simos et al., 1999) | English | Object picture naming | Color drawings of simple objects |
| (Hamberger, Goodman, Perrine, & Tamny, 2001) | English[2] | Object picture naming | Snodgrass and Vanderwart (1980) |
| (Ojemann, Ojemann, & Lettich, 2002) | English (1 Korean) | Object picture naming | Black and white drawings of common objects |
| | | Verb generation from a noun | Concrete nouns |
| (Roux & Trémoulet, 2002) | French (and participants' L2) | Object picture naming → "This is…" | |
| (Roux et al., 2003) | French | Object picture naming<br>Verb generation from a picture | Images representing concrete nouns |
| (Lubrano, Roux, & Démonet, 2004) | French | Object picture naming → "This is…" | |
| (Lucas, McKhann, & Ojemann, 2004) | English (and participants' L2) | Object picture naming (in 2 languages) | Line drawings of common objects |
| (Walker, Quiñones-Hinojosa, Berger, & Grossman, 2004) | English (and participants' L2) | Object picture naming (in 2 languages) | Line drawings of common objects |
| (Roux, Lauwers-Cances, Trémoulet, Mascott, & Démonet, 2004) | French (and participants' L2) | Object picture naming → "This is…" | |
| (Corina et al., 2005) | English | Object picture naming<br>Verb action naming → "-ing" | Vignettes of actors carrying out common actions |
| (Bello & Acerbi, 2006) | Multilingual | Object picture naming<br>Verb action naming | Batteria per l'analisi dei déficit afasici (BADA) (Miceli, Capasso, Laudanna, & Burani, 1994) |

[1] The language was not explicitly stated in the text. We inferred it from the team location and/or other published papers by the same author(s).

| Study | Language | Tasks | Stimuli |
|---|---|---|---|
| (Sato et al., 2006) | Japanese | Object picture naming | Common objects |
| (Petrovich Brennan et al., 2007) | English | Object picture naming | Drawings from BDAE and Snodgrass and Vanderwart (1980) |
| (Ilmberger et al., 2008) | German | Object picture naming → "This is…" | Line drawings |
| (Mandonnet, Gatignol, & Duffau, 2009) | French | Object picture naming | DO 80 |
| (Moritz-Gasser & Duffau, 2009) | French | Object picture naming → "This is…" | DO 80 |
| (Roux, Borsa, & Démonet, 2009) | French | Object picture naming → "This is…" | |
| (Roux, Boukhatem, Draper, Sacko, & Démonet, 2009) | French | Object picture naming → "This is…" | |
| (Hamberger, Seidel, Goodman, & McKhann, 2010) | English | Object picture naming → "This is…" | Line drawings of common objects |
| (Cervenka, Boatman-Reich, Ward, Franaszczuk, & Crone, 2011) | (Participants' L1) and English | Object picture naming | BNT |
| (Giussani et al., 2011) | French[1] | Object picture naming | Living and non-living, controlled for frequency |
| (Papagno et al., 2011) | Italian[1] | Object picture naming<br>Verb action naming | Living and non-living objects and actions |
| (Papagno et al., 2011) | Italian | Object picture naming<br>Verb generation from a picture | Living and non-living objects and actions |
| (Gil-Robles et al., 2013) | Spanish[1] | Object picture naming | DO 80 |
| (Moritz-Gasser, Herbet, & Duffau, 2013) | French | Object picture naming | DO 80 |
| (Tarapore et al., 2013) | English[1] | Object picture naming | Snodgrass and Vanderwart (1980) |
| (Conner, Chen, Pieters, & Tandon, 2014) | English | Object picture naming<br>Verb action naming → "-ing" | Line drawings |

| Study | Language | Tasks | Stimuli |
| --- | --- | --- | --- |
| (Khan, Herbet, Moritz-Gasser, & Duffau, 2014) | French | Object picture naming → "This is…" | DO 80 |
| (Krieg et al., 2014) | German and English | Object picture naming → "This is…" | Black and white drawings of common objects |
| (Lubrano, Filleron, Démonet, & Roux, 2014) | French | Object picture naming | Center for Research in Language-International Picture Naming Project corpus CRL-IPNP (Székely et al., 2005) |
| | | Verb action naming | |
| (Tomasino et al., 2014) | Serbian and Italian | Object picture naming<br>Verb generation from a picture | Snodgrass and Vanderwart (1980) |
| (Havas et al., 2015) | Spanish | Object picture naming<br>Verb action naming | Line drawings of objects and actions controlled for linguistic variables |
| (Ille, Sollmann, Hauck, Maurer, Tanigawa, Obermueller, Negwer, Droese, Zimmer, et al., 2015) | German | Object picture naming → "This is…" | Color pictures of common objects |
| (Ille, Sollmann, Hauck, Maurer, Tanigawa, Obermueller, Negwer, Droese, Boeckh-Behrens, et al., 2015) | German[1] | Object picture naming | Color pictures of common objects |
| (Li et al., 2015) | Chinese | Object picture naming | |
| (Rofes, Spena, Miozzo, Fontanella, & Miceli, 2015) | Italian | Object picture naming<br>Finite verb production | (Rofes, de Aguiar, & Miceli, 2015) |
| (Roux et al., 2015) | French | Object picture naming | Snodgrass and Vanderwart (1980) |
| (Rutten, 2015) | Dutch[1] | Object picture naming → "This is…" | |
| (Sierpowska et al., 2015) | Spanish | Object picture naming<br>Noun based verb generation | |
| (Hamberger et al., 2016) | English | Object picture naming | Pictures of common objects |

| Study | Language | Tasks | Stimuli |
|---|---|---|---|
| (Riva, Casarotti, Comi, Pessina, & Bello, 2016) | Italian | Object picture naming | |
| (Riva, Fava, et al., 2016) | Italian[1] | Object picture naming | |
| (Skrap, Marin, Ius, Fabbro, & Tomasino, 2016) | Italian | Object picture naming<br>Verb naming | Selected from batteries available in the Italian normative data. |
| (Sollmann et al., 2016) | German[1] | Object picture naming | |
| (Wongsripuemtet et al., 2016) | Thai | Object picture naming | |
| (Rofes et al., 2017) | Italian | Object picture naming → "This is…"<br>Finite verb production | (Rofes, de Aguiar, et al., 2015) |
| (Roux, Durand, Djidjeli, Moyse, & Giussani, 2017) | French | Object picture naming | Snodgrass and Vanderwart (1980) |
| (Chang et al., 2018) | Chinese | Object picture naming | DO 80 |
| (Ferpozzi et al., 2018) | Italian[1] | Object picture naming | |
| (Forseth et al., 2018) | English | Object picture naming | Drawings from BDAE and Snodgrass and Vanderwart (1980) |
| (Herbet, Moritz-gasser, Lemaitre, & Duffau, 2018) | French | Object picture naming | DO 80 |
